# Motor cortex latent dynamics encode spatial and temporal arm movement parameters independently

**DOI:** 10.1101/2023.05.26.542452

**Authors:** Andrea Colins Rodriguez, Matthew G. Perich, Lee Miller, Mark D. Humphries

## Abstract

The fluid movement of an arm requires multiple spatiotemporal parameters to be set independently. Recent studies have argued that arm movements are generated by the collective dynamics of neurons in motor cortex. An untested prediction of this hypothesis is that independent parameters of movement must map to independent components of the neural dynamics. Using a task where monkeys made a sequence of reaching movements to randomly placed targets, we show that the spatial and temporal parameters of arm movements are independently encoded in the low-dimensional trajectories of population activity in motor cortex: Each movement’s direction corresponds to a fixed neural trajectory through neural state space and its speed to how quickly that trajectory is traversed. Recurrent neural network models show this coding allows independent control over the spatial and temporal parameters of movement by separate network parameters. Our results support a key prediction of the dynamical systems view of motor cortex, but also argue that not all parameters of movement are defined by different trajectories of population activity.

**Significance Statement:** From delicate strokes while drawing to ballistic swings while playing tennis, a skilled arm movement requires precise control of both its direction and speed. Motor cortex is thought to play a key role in controlling both, but it is unclear how they are jointly controlled. We show here that the population activity in motor cortex represents both the spatial and temporal properties of arm movements in the same low-dimensional signal. This representation was remarkably simple: the movement’s direction is represented by the trajectory that signal takes; the movement’s speed by how quickly the signal moves along its trajectory. Our network modelling shows this encoding allows an arm movement’s direction and speed to be simultaneously and independently controlled.

## Introduction

Motor cortex is increasingly viewed as a dynamical system, whose low-dimensional dynamics generate movement (Churchland et al., 2010, 2012; Shenoy et al., 2013; Michaels et al., 2016; Gallego et al., 2017, 2018; Russo et al., 2018; Perich et al., 2018b; Even-Chen et al., 2019; Vyas et al., 2020). This perspective moves from a simple “representational” view that single neuron activity encodes movement parameters to posit that much of the activity in motor cortex reflects the ongoing evolution of a dynamical system, which results in a descending movement execution signal (Shenoy et al., 2013; Kaufman et al., 2014; Gallego et al., 2018; Russo et al., 2018). It has been used to account for the variety of complex single neuron responses during simple movements (Churchland et al., 2012; Hennequin et al., 2014) and for changes in neuron tuning between preparation and execution (Churchland et al., 2010; Elsayed et al., 2016). But if the low-dimensional dynamics in motor cortex are to generate movement, then the question of representation remains: of how changes in those dynamics map to changes in the parameters of movement (Russo et al., 2018; Humphries, 2021a; Saxena et al., 2022).

Arm movements are well-suited to addressing this question as they have parameters that can be independently specified (Gordon et al., 1994; Favilla and De Cecco, 1996; Vindras et al., 2005). An arm movement has spatial parameters, its direction and extent, and temporal parameters, including its speed. Subjects reaching towards a fixed target can readily adapt how quickly they move their arm given cues or explicit instructions (Soechting and Lacquaniti, 1981; Nishikawa et al., 1999; Mazzoni et al., 2007). This ability to specify the spatial and temporal parameters of arm movement independently predicts that, if the low-dimensional dynamics of motor cortex generate arm movements, then changes in those dynamics should independently specify these spatial and temporal parameters. Yet little is known about how neural dynamics can specify these parameters simultaneously.

Here we focus on testing this prediction by studying how motor cortex simultaneously encodes both the direction of a reaching movement and how quickly that movement is made. There has been considerable study of how neural activity in motor cortex correlates with either of these spatial or temporal parameters of arm movement, but not both. Changes in the activity of individual neurons in motor cortex correlate with the direction of an arm movement (Georgopoulos et al., 1982, 1986; Ashe and Georgopoulos, 1994; Moran and Schwartz, 1999) or with how quickly the arm is moved (Schwartz, 1992, 1994; Moran and Schwartz, 1999; Paninski et al., 2004; Churchland et al., 2006; Wang et al., 2007; Inoue et al., 2018). Researchers taking a dynamical systems perspective on motor cortex have observed that changes in the trajectory of its population activity through state-space correlate with different directions of arm movement (Santhanam et al., 2009; Churchland et al., 2012; Michaels et al., 2016; Russo et al., 2018; Even-Chen et al., 2019) and, more recently, with the speed of arm movement (Saxena et al., 2022; Schroeder et al., 2022). However, most studies examine movement parameters one at a time, leaving unaddressed how independent parameters of arm movement map to independent dynamics of the motor cortex.

To address this question, we analysed population activity in motor cortex from monkeys performing a self-paced sequential-target task that extensively sampled variations in both the direction and speed of planar arm movements. We report here that, as predicted from behaviour, direction and speed are independently encoded within the low dimensional trajectory of population activity. We find that how quickly the arm moves is encoded by how quickly a fixed trajectory of population activity is traversed, not by changes in the trajectory itself, disagreeing with recent results on speed encoding in motor cortex (Sax-ena et al., 2022). We then use recurrent neural network modelling to show this encoding allows the rate of traversing a neural trajectory to be controlled independently from the trajectory itself.

## Methods and Materials

### Subjects and task

We analysed 13 population recordings from the motor cortex (6 from PMd and 7 from M1) of three monkeys (Monkey M, T and C). Monkey C contributed 3 populations from M1, Monkey T contributed 3 populations from PMd and Monkey M, which had implants in M1 and PMd simultaneously, contributed 6 populations. This gave a total of 10 behavioural sessions. Data from 4 populations (monkey M and T, PMd and M1 areas) are publicly available (Perich et al., 2018a).

Monkeys were trained to reach four targets that sequentially appeared on the screen, using a cursor controlled by a planar manipulandum. Hand movements were constrained to the horizontal plane on a workspace of 20 × 20 cm. Each target’s position was chosen by a pseudo-random algorithm which selected the distance (5-15 cm) and angle (0-360*^◦^*) with respect to the previous target.

Targets appeared 196 ms after the monkey reached the previous target, but only if they stayed on the previous target for at least 100 ms. The target acceptance window was a 2 cm square centred on the target position.

We define movement direction for the first reach of the sequence as the direction between the hand position at the moment of target appearance and the first target’s position. For later reaches, the movement direction corresponds to the direction between the positions of the current and next targets.

Only successful trials were used for the following analyses to ensure consistency across reaches between targets. Failed trials by, for example, leaving the centre of the workspace or not holding the target for the required minimum time, were discarded.

All surgical and experimental procedures were consistent with the guide for the care and use of laboratory animals and approved by the institutional animal care and use committee of Northwestern University.

### Neural recordings

Subjects were implanted with 100-electrode arrays (Blackrock Microsystems, Salt Lake City, UT) in PMd (1 mm electrode shaft length) and M1 (1.5mm length). Spike sorting was performed manually using standard techniques. See (Lawlor et al., 2018; Perich et al., 2018a; Glaser et al., 2018) for a detailed description of the neural recordings. Each population recording contains between 24-95 units.

### Data processing

Spike trains were filtered with a Gaussian filter of standard deviation *σ*. For each population, we defined *σ* as the median of its inter-spike interval (*σ* = 36 −> 86 ms for all populations). We aligned the neural activity to movement onset defined by a hand-movement speed threshold of 8 cm/s (Lawlor et al., 2018). The movement’s end corresponded to the time when the hand entered the acceptance window. The movement duration was defined as the time between the movement’s onset and end.

Arm movements were binned according to their direction (8 bins, as in Fig. 1) and duration (4 bins, 200-300, 300-400, 400-500 and 500-600 ms), resulting in 32 conditions. To make all neural activity in a duration bin have equal length, we rounded the duration of the movement to the largest end of their duration bin. For example, if the associated movement lasted 240 ms, then the movement’s end was considered to be 300 ms after movement onset. Because all movements were self-paced, reaction times varied across movements. To ensure that all times that the population encoded the current movement were selected, for each movement we selected the corresponding neural activity from 500 ms before movement onset until 300 ms after the movement ended (600-900 ms from movement onset) and averaged the neural activity across movements.

**Figure 1:**
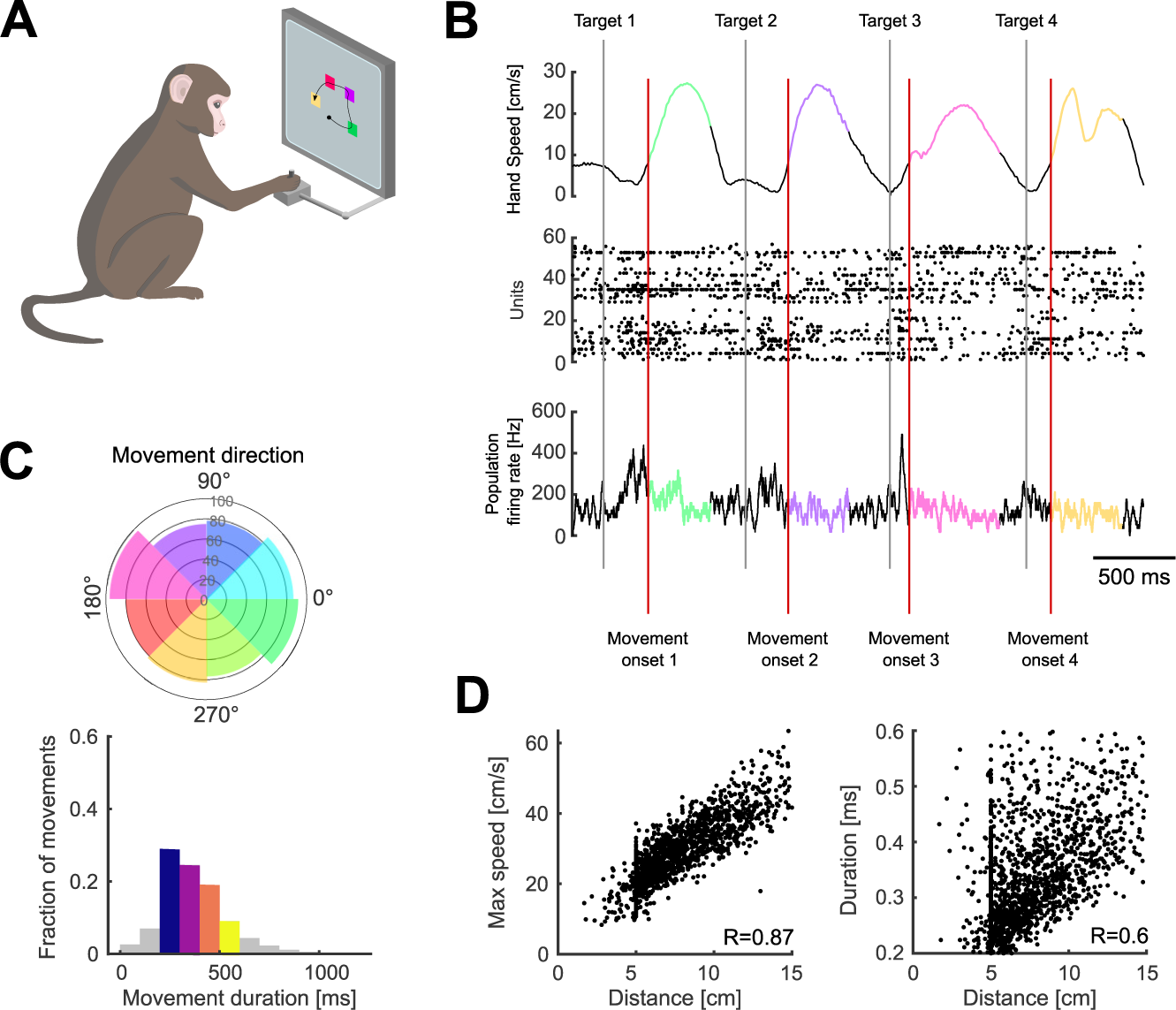
Variability of arm movement behaviour in a sequential-target task. A) Task schematic. Monkeys were trained to reach four targets that appeared sequentially on the screen, using a planar manipulandum (image adapted from (Colins Rodriguez, 2021)). B) Hand speed (top), spike rasters (middle), and firing rate (bottom) for an example trial. Grey and red vertical lines indicate the times of target appearance and movement onset, respectively. Highlighted segments show movement execution coloured by movement direction. C) Variability of arm movement parameters for an example session: movement direction (top; radius shows the number of movements), duration (bottom). Coloured direction slices show the bins used for direction-based analyses. Coloured bars for duration indicate the bins used for duration-based analyses. D) Correlation between maximum arm movement speed and distance (left) and duration and distance (right) for one example session.

### Finding neural trajectories

We performed Principal Component Analysis (PCA) and demixed Principal Component Analysis (dPCA) (Kobak et al., 2016) across all conditions’ averaged neural activity. For both methods, neural activity was preprocessed in the same way: Neurons that showed less than 5 spikes across all conditions were discarded; and because the variance of each neuron varies with its firing rate, we balanced the variance across the population by soft-normalising the responses of each neuron (normalisation factor= firing rate range + 5) before applying PCA (Russo et al., 2020; Saxena et al., 2022).

We used the published Matlab package of dPCA (https://github.com/machenslab/dPCA) to test if movement parameters were encoded in specific subspaces. Considering our parameters of interest, the following marginalisations were used: direction, duration and condition-invariant. To equalise the length of the neural activity, the average neural activity of each condition was temporally scaled to 600 time bins using linear interpolation before performing dPCA. The total number of dimensions extracted was set to 15 for all populations. The neural trajectories of each component (Fig. 2B-D) were obtained by projecting the neural activity onto each direction-, duration- and condition-independent component. The variance explained by a parameter (e.g. direction) was then the sum of the variances explained by the dPCs corresponding to that parameter (Fig. 2E).

**Figure 2:**
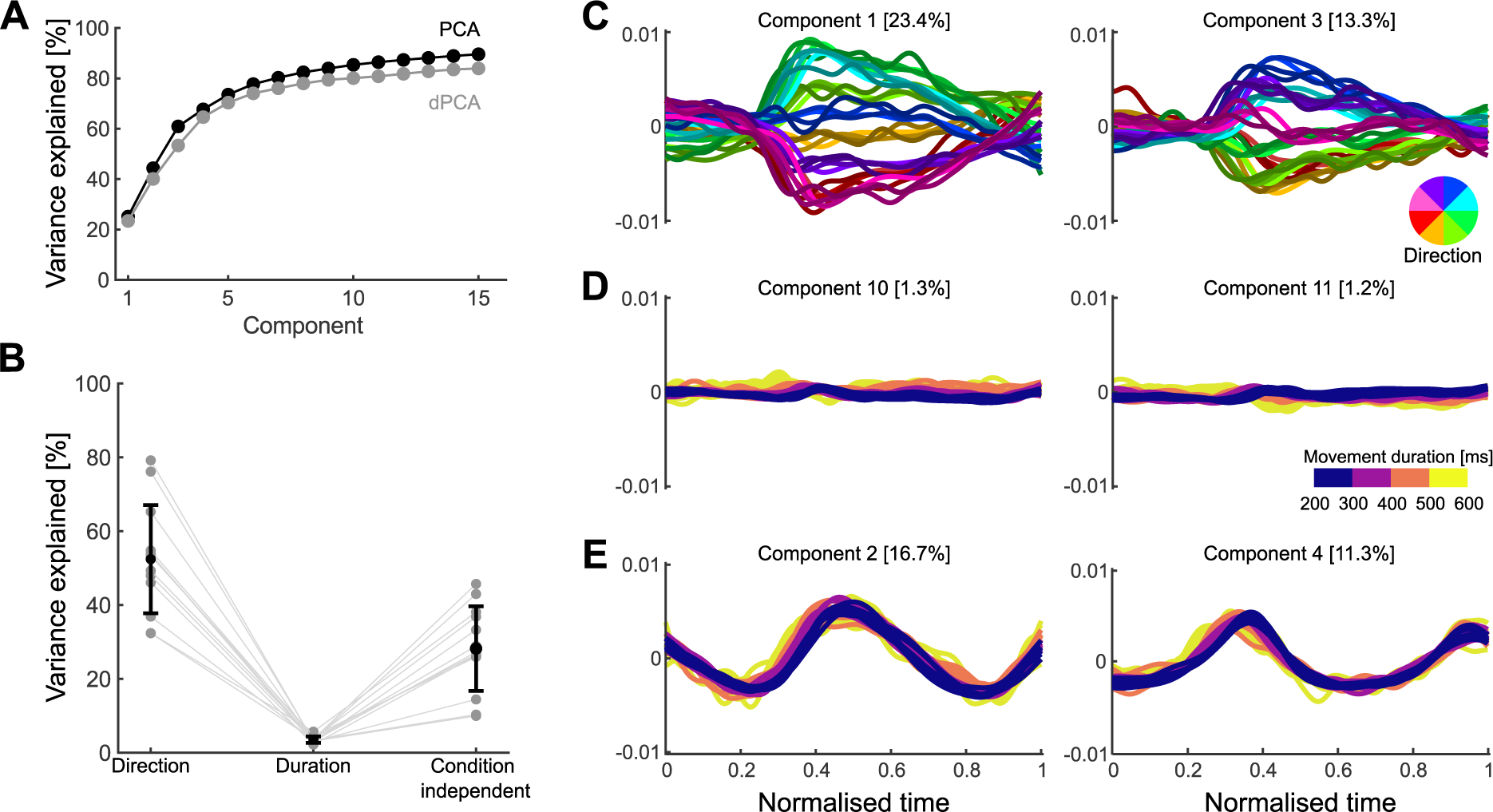
No encoding subspace for how quickly a movement was made. A) Variance of neural activity explained as a function of the number of retained principal components for PCA and dPCA. B) Variance explained by the direction, duration, and condition-independent components. Grey dots show the total for each population, black dots the average across populations, and error bar the standard deviation. C) Population activity projected onto the first two direction components. Trajectories, one per condition, are coloured by movement direction and darker shades indicate longer durations. D) Population activity projected onto the first two duration components. Trajectories, one per condition, are coloured according to movement duration. E) Population activity projected onto the condition-independent components.

For PCA analyses we used the same preprocessing as for dPCA except that neural activity was not temporally scaled. We used PCA to define a common subspace (for direction and duration) for each population. The embedding dimensions *D* of each common subspace were chosen as the top *D* Principal Components (PCs) necessary to explain 80% of the variance of the neural activity. We obtained the neural trajectories by projecting the population neural activity onto the common subspace.

### Comparing time-scaled trajectories

We temporally scaled the neural trajectories by linear interpolation of their values (Fig. 4B,C). All trajectories were scaled from their original duration to a final size of 600 time bins. To quantify how well two temporally scaled trajectories matched (Fig. 4C), we computed their coefficient of determination (*R*^2^). For this, we collapsed the temporally scaled low-dimensional trajectories ***X*** (composed of points ***x*** of *D* dimensions) and ***Y*** (composed of points ***y*** of *D* dimensions) each into a single vector containing all dimensions concatenated (length ***x*** *· D*). We then computed the coefficient of determination

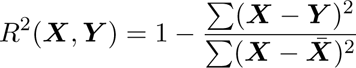

Where 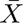 denotes the mean of *X*. Therefore, *R*^2^=1 indicates perfect matching of the trajectories, *R*^2^=0 indicates trajectories match as well as the mean value of the trajectories (baseline) and *R*^2^ *<* 0 indicates that matching of the trajectories is lower than this baseline model, meaning they are of different magnitudes.

To compare how similar two trajectories corresponding to the same duration and direction bin could be, we randomly divided all the movements corresponding to the same direction and duration bin (within-bin) into two equal-sized sets. We then created two trajectories by taking the average neural activity of each set and projecting them into the low-dimensional subspace. The coefficient of determination was calculated between these two trajectories. We repeated these three steps for half the number of movements corresponding to each duration and direction bin. We then pooled all the values across directions and durations to compute the average for each population (Fig. 4C).

### Comparing distances between trajectories

To compare the distance between trajectories of different lengths, we used the Hausdorff distance, the maximum of the minimum distance between two sets of points (Fig. 4D-F). The Hausdorff distance between two trajectories ***X*** (composed of points ***x***) and ***Y*** (composed of points ***y***) is defined as

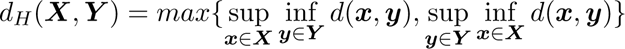

where *d* quantifies the Euclidean distance between ***x*** and ***y***, sup represents the supremum and inf the infimum.

To compare the Hausdorff distance between trajectories corresponding to different durations, we measured the Hausdorff distance between each trajectory and all other trajectories corresponding to the same direction and different durations (3 other duration bins) (Fig. 4D).

To compare the Hausdorff distance between trajectories corresponding to different directions, we measured the distances between each trajectory and the trajectories corresponding to the 2 adjacent direction bins (one to the left and one right) that had the same duration. To test if the distances between trajectories corresponding to different durations were lower than trajectories corresponding to different directions, a *t*-test was performed for each population (*N* = combinations of duration bins (3)× adjacent bins (2)× Durations (4) × Directions (8)=192).

To compare the Hausdorff distance between trajectories corresponding to different movements’ speeds (Fig. 4F), we performed similar analyses. Because movement distance and speed were highly correlated (Fig. 1D), we selected movements that only were within a narrow window of distances (4-6 cm). We then divided these movements into high and low speeds and according to their direction (8 bins, 16 conditions total). We used the median speed in each dataset as a threshold to classify the movements. We scaled the neural activity of all trials to a reference duration and took the average for each condition. We projected the average neural activity of these 16 conditions into the common subspace that encompassed direction and duration. Similar results were obtained by first defining a speed-direction subspace using PCA on the average neural activity for the 16 conditions, and then projecting the per-condition activity into that subspace. To compare the distance between trajectories we followed the procedure described above for different durations and directions. Specifically, we compared the distance between high and low speed trajectories versus the distance between trajectories corresponding to adjacent direction bins (Fig. 4F, left).

To test if distances between trajectories corresponding to different speeds were significantly lower than the distances between trajectories corresponding to different directions, a *t*-test was performed for each population (*N* = Speed bins (2) × Adjacent bins (2) × Directions (8)=32).

### Decoding movement duration

To predict a movement’s duration from neural data, we built a linear decoder that used as input a reference neural trajectory that belongs to the same direction bin as the sample and whose duration is known (Figure 5).

Since the reference and the sample trajectory are hypothesised to share their geometry, for each point in the sample *S*(*t_i_*) we can find the closest point in the reference trajectory *R*(*idx_i_*), where the time index *idx_i_*is calculated as:

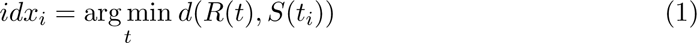

where *d* denotes the Euclidean distance (Remington et al., 2018). The relative speed (*v*) between the reference and sample trajectories was calculated as the slope of a linear regression between the times of the sample *t*_1_*_,…,n_* and the *idx_i_* values. Therefore, the duration of a sample trajectory can be predicted by the linear decoder: *Duration_sample_*= *vDuration_ref_*. Because the neural trajectories were selected from the start to the end of movement encoding, the movement duration could be directly computed from the duration of the entire trajectory (*Duration_sample_*) by subtracting the time outside the movement execution (500 ms for PMd and 450 for M1). For all populations, the 8 trajectories that corresponded to movements lasting between 300 and 400 ms were selected as the references. The remaining 24 trajectories were used as samples for which durations were predicted.

### RNN modelling

To examine how a population of neurons could control both the geometry of a neural trajectory and how quickly it is traversed, we implemented a recurrent neural network.

Since we observed the same key encoding features in PMd and M1, we implemented a single neural network that reproduced the separation and temporal scaling of the trajectories. Further work could aim to reproduce the differences found between these two regions (such as the difference in timing in Fig. 3) and analyse how they communicate.

**Figure 3:**
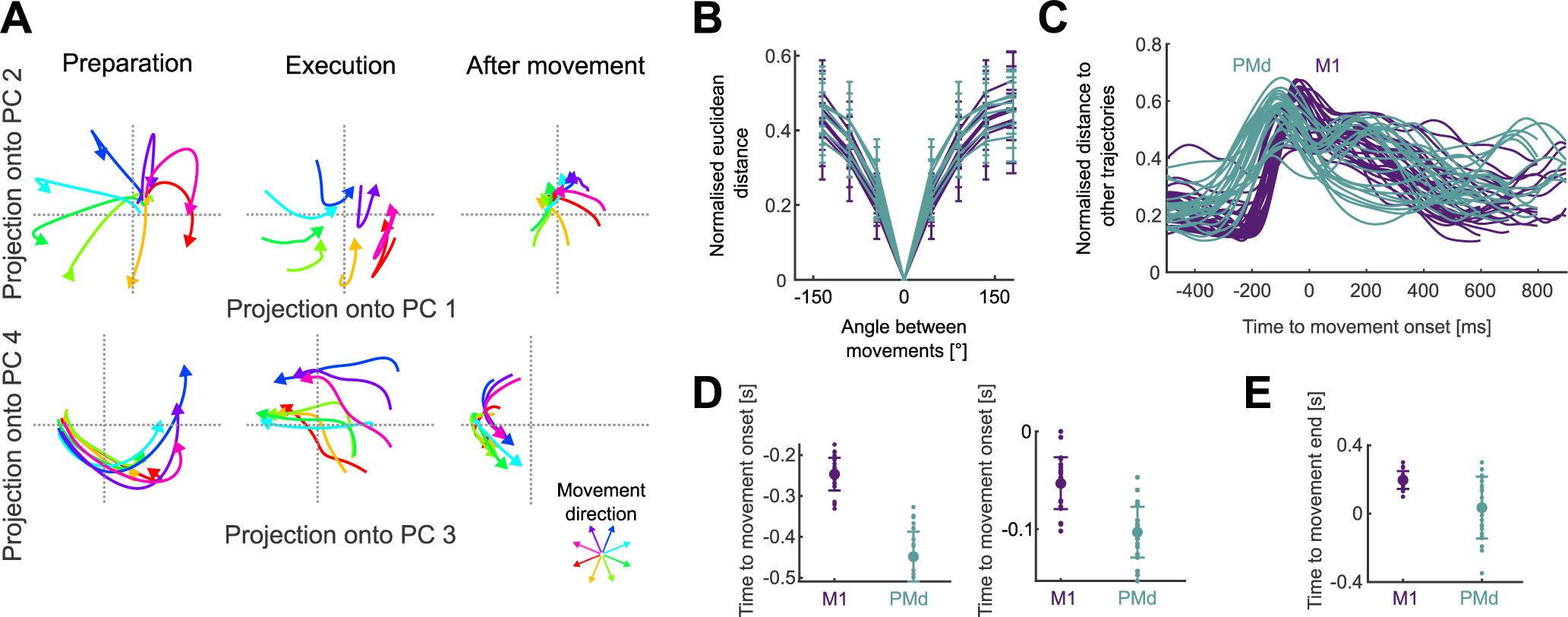
Neural trajectories encode arm movement direction. A) Direction-dependent neural trajectories of movements between 200 and 300 ms duration from an example M1 population. Neural trajectories were taken from 250 ms before movement onset until 200 ms after movement end and divided here into 3 stages: preparation, the time up to movement onset (left); execution, the time between movement onset and movement end (middle); and after movement, from when the hand reached the target (right). B) Mean distance between neural trajectories as a function of the difference in the direction of their corresponding arm movements (of 200-300 ms duration). One line per population. Error bars indicate SD. Distances were normalised by the maximum distance between the trajectories of each population. C) Change in the distance between neural trajectories over time, averaged over all movement directions of the same duration (N=52 lines: 4 duration bins × 13 populations). D) Times of minimum distance between trajectories before movement onset (left) and of maximum distance between trajectories (right). E) Time of minimum distance between trajectories after movement onset.

We based our RNN on the stability-optimised circuit model of Hennequin et al. (2014), a rate-based network (*N* = 200 units, 100 excitatory, 100 inhibitory) whose temporal evolution is described by:

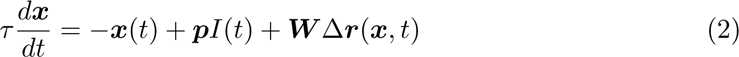

where ***x*** is the vector of potentials, *τ* = 200 ms is the network’s time constant, *I*(*t*) is the ramping input signal, ***p*** is the vector defining the projection of the input to the units, ***W*** represents the connectivity matrix and Δ***r*** contains the instantaneous firing rate of all units with respect to a baseline.

We implemented the stability optimisation algorithm for ***W*** detailed in (Hennequin et al., 2014), creating 20 realisations of ***W*** for which we obtained similar results to those shown for the example network in Fig. 6.

The instantaneous firing rate Δ***r*** was defined as follows:

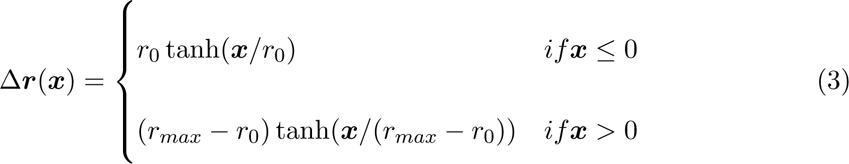

with baseline firing rate *r*_0_= 5 Hz and maximum firing rate *r_max_*= 70 Hz.

For all of our simulations, following Hennequin et al. (2014) we used a 1 s ramping exponential input *I*(*t*), whose time constant was fixed (200 ms), and a subsequent 200 ms decay (time constant= 0.1 ms). In our simulations, the start of the input (*t*= 1 s) is equivalent to the beginning of movement preparation, while the return to a steady-state corresponds to a time after movement ends. All simulations lasted 4 s. The strength of the input projection onto each unit was drawn from a uniform distribution between [−>1, 1]. We also tested inputs drawn from a uniform distribution between [0, 1] to model solely excitatory inputs to the cortical circuit and found similar results.

To simulate the effect of different directions, we sampled one vector of input projection strengths ***p*** and rotated it 10 times to span 360*^◦^* so that each input rotation putatively corresponded to a different direction of arm movement. Each rotation of the *n*-dimensional vector ***p*** was achieved by independently rotating each 2D plane taken sequentially from the *n*-dimensional vector. We then defined the network’s output subspace by using PCA across the 10 resulting time-series of network activity.

To test control of how quickly a trajectory is traversed we fixed the input projection vector ***p*** but varied the network’s time constant *τ* = [160, 200, 240] ms. We projected the resulting neural activity onto the network’s output subspace. We defined the duration of each trajectory as the time it took for the trajectory to return to the region of baseline pre-input activity, which was defined as a hypersphere centred on the average start of the trajectories and radius equal to the 20*^th^* percentile of all distances between points in each population.

## Results

Three monkeys were trained to move a cursor to targets on a screen using a planar manipulandum (Fig. 1A). In each trial, four targets appeared sequentially, each new target appearing in a pseudo-random position after the monkey reached the previous target and held for 200 ms (Fig. 1B). Successfully hitting all four targets resulted in a liquid reward at the end of each trial. We consider henceforth each movement within a successful sequence of four to be an individual arm movement. Extracellular neural activity from dorsal premotor cortex (PMd) and/or primary motor cortex (M1) was recorded during the task using “Utah” arrays (Blackrock Neurotech, Salt Lake City). We analyse here 6 PMd and 7 M1 population recordings from 10 sessions, each population containing between 24 and 95 neurons.

### Task-induced variation in independent arm movement parameters

This task design induced variability in both the direction of arm movements and how quickly they were made (Fig. 1C). The maximum speed of movement was strongly correlated with the distance to the target (Fig. 1D, mean *R* = 0.8 across populations). Because of this strong scaling of maximum arm speed with distance, the duration of each movement was less correlated with the distance to the target (mean *R* = 0.56, Fig. 1D) and with the maximum arm speed (mean *R* = 0.27). As we sought to study how the spatial and temporal parameters of arm movement were independently encoded, we thus use movement duration here to capture how the profile of instantaneous hand movement speed varied from movement to movement (examples in Fig. 1B), though we do not mean to suggest that this summary measure of the temporal parameter of movement is directly specified by motor cortex.

### How quickly a movement is made is not encoded in a subspace of activity

One way to simultaneously encode multiple parameters of movement is to represent them in distinct subspaces of the population activity (Kaufman et al., 2014; Kobak et al., 2016). To test this possibility, we performed demixed Principal Component Analyses (dPCA) (Kobak et al., 2016). dPCA defines the subspaces that best represent chosen task-related variables while maximising the captured variance of the original neural activity. In this way, dPCA creates a low-dimensional representation of the neural activity in a coordinate system where each axis is associated with a parameter of interest.

We looked for subspaces in the neural activity that separately encoded movement direction and how quickly a movement is made. We binned the movements into eight directions and four durations, as shown in Fig. 1C, giving 32 conditions in total, normalised the condition-averaged neural activity from movements of different durations to a common time-base, and applied dPCA (Methods).

For all populations, dPCA captured almost the same total variance as did standard Principal Component Analyses (PCA) (Fig 2A). The direction and condition-independent components consistently captured the majority of the variance (Fig 2B, averages of 52% and 28% respectively), while the duration components captured only 3.5% on average (Fig 2B). While projections of neural activity on to the direction components were clearly separated over time (Fig 2C), projections on to the duration components had virtually no separation (Fig 2D). These results suggest that the direction of arm movement was encoded by changes of neural activity within a defined subspace, but how quickly it moved was not.

### Arm movement direction is encoded in the trajectory of population activity

Because there were not distinct subspaces of activity for the direction of arm movement and for how quickly it moved, we took an unsupervised approach to examine how these independent parameters were simultaneously encoded in the neural dynamics. For each population, we found the average neural activity for each of the 32 conditions, then performed PCA across all conditions to define a common subspace. In contrast to dPCA, PCA does not require equal length neural activity and so we did not need to rescale time. We defined the size of the subspace as the top *D* dimensions required to explain at least 80% of the variance of the neural activity; this ranged from 3-11 dimensions across populations. Projecting a population’s activity into its PCA-defined subspace allowed us to define its trajectory through neural state-space over time (Fig. 3A shows an example in pairs of dimensions; all subsequent analyses of a population’s trajectories were performed in its full *D*-dimensional subspace). We then sought to quantify how these trajectories encoded both the direction and speed of arm movement. We first focus on the encoding of direction.

Across all populations, we observed that the neural trajectories started in the same location in state-space, separated for different directions of movement, then reconverged after the movement ended (Fig. 3A). The separation of trajectories in both M1 and PMd was monotonically proportional to the difference in arm movement direction (Fig. 3B), consistent with the idea that movement direction is encoded by the trajectory of neural activity.

We then asked when the encoding of direction started and ended in the neural trajectories. To estimate this, we determined when the trajectories corresponding to different movement directions first separated and then reconverged. We found the separation and reconvergence of trajectories in M1 consistently lagged those in PMd (Fig. 3C). Trajectories in PMd reached a minimum separation before movement onset about *∼*200 ms earlier than those in M1 (Fig. 3D, left), in agreement with previous studies (Thura and Cisek, 2014). The maximum separation of trajectories also happened first in PMd (Fig. 3D, right). After movement onset, PMd trajectories reconverged earlier than those in M1 (Fig. 3E). Direction coding by the separation of neural trajectories thus started and ended earlier in PMd than it did in M1.

To compare the trajectories in PMd and M1 in subsequent analyses, we took their neural trajectories between when they first separated (Fig. 3D, left) and then reconverged (Fig. 3E). These were 450 ms before movement onset to 50 ms after the end of movement for PMd, and 250 ms before movement onset to 200 ms after the end of movement for M1. Both PMd and M1 trajectories thus showed encoding of movement direction that started before movement onset and was sustained beyond the end of arm movement.

### How quickly the arm moves is encoded in the timing of the neural trajectories

We then asked how the neural trajectories could also encode how quickly the arm moves. We considered two hypotheses. First, the *geometry* hypothesis: as with direction, movement speed is represented by different trajectories through neural state space, with slower movements generated by longer trajectories (Fig. 4A, left). Alternatively, the *temporal scaling* hypothesis: the trajectory through neural state space is fixed for a given direction of movement, but simply traversed at a different rate, with slower movements generated by slower traversal of trajectories (Fig. 4A, right).

**Figure 4:**
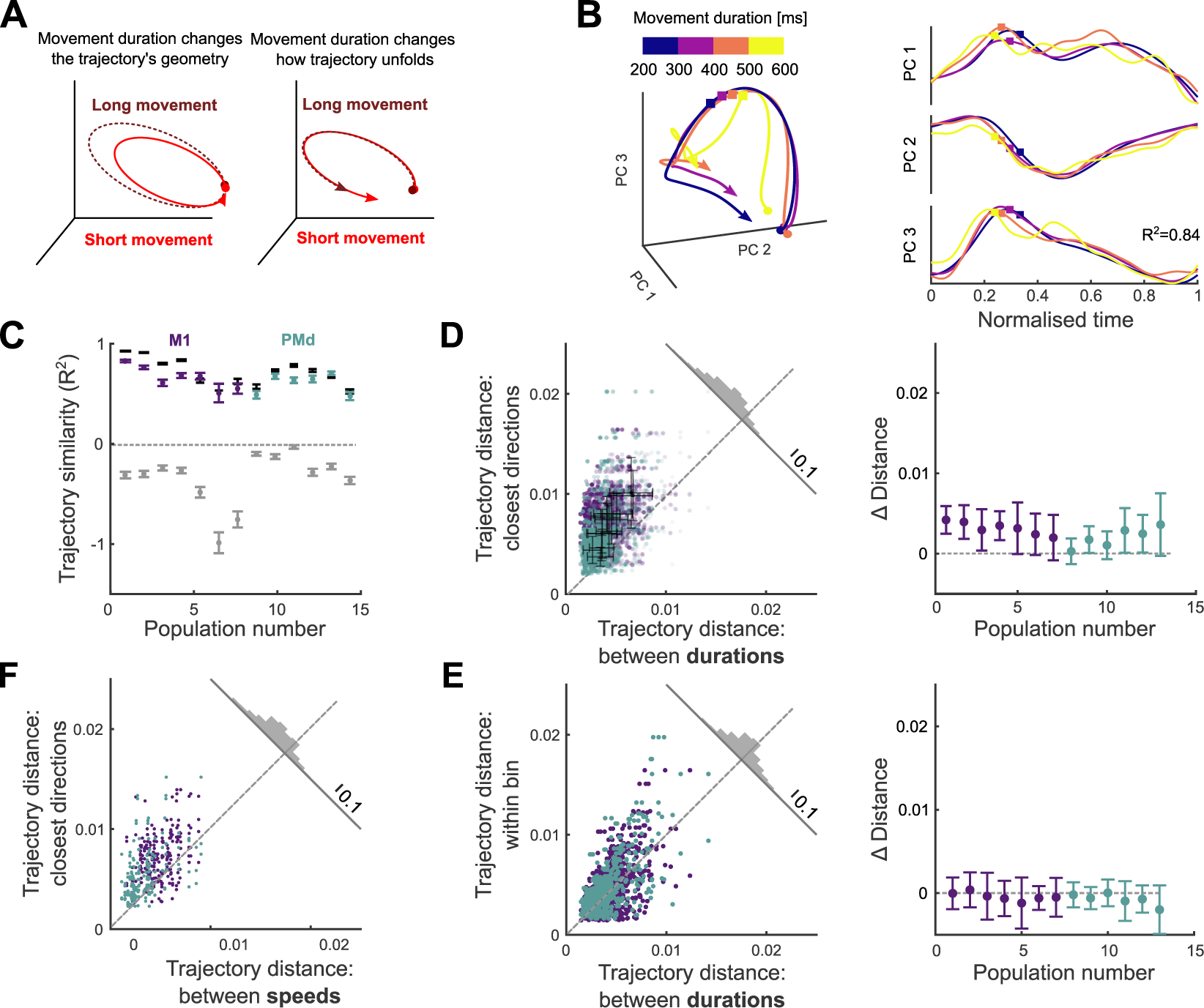
How quickly the arm moves is encoded by the traversal of fixed neural trajectories. A) Hypotheses for how neural trajectories may encode how quickly the arm moves: in changes of the trajectory itself (left) or in how the trajectory unfolds over time (right). B) Left: Projection of neural trajectories prior to temporal scaling, corresponding to a given direction but different movement durations, each plotted for 600 ms. Coloured circles and squares mark start of the preparation and movement onset, respectively. Right: Trajectories from the left panel normalised by their duration. *t* = 0 is the time of minimal separation between trajectories and *t* = 1 is the return to that baseline. Coloured squares mark movement onset. C) Similarity of time-scaled trajectories corresponding to the same direction (purple/teal bars), measured by the coefficient of determination *R*^2^. Grey bars are lower limits for *R*^2^ from comparing neural trajectories for movements of the same duration but in different directions. Black bars give the estimated upper limit for *R*^2^ from sub-sampled trajectories from the same duration and direction. *N* between 171 and 1165 per population. Error bars are SEM. D) Hausdorff distance between trajectories from adjacent direction bins compared to the Hausdorff distance between trajectories corresponding to different durations of movements in the same direction. Opacity of each data point is proportional to the number of samples used to produce each trajectory. Solid colour shows maximum number of samples. Black error bars indicate mean and SD for each population. Histogram shows the distribution over all data-points of the distance to the dashed line. Right: Mean difference for each population between the *x*- and *y*-axis values of the left panel. E) Hausdorff distance between trajectories of the same direction and duration compared to the Hausdorff distance between trajectories corresponding to different durations of movements in the same direction (*x* axis in panel D, left). Right: Difference between the *y* and *x* axes of the left panel. F) Same as D) left, for trajectories corresponding to different speeds.

Results from the dPCA analysis in Figure 2 suggested there was no subspace of neural activity related to movement duration, providing initial evidence against the geometry hypothesis. Here we further test the two hypotheses in the unsupervised subspace we obtained with PCA.

We observed that neural trajectories corresponding to the same direction yet different durations of arm movement closely overlap each other in state space (Figure 4B, left). Following temporal rescaling, these trajectories were strikingly similar over time (Fig. 4B, right). To quantify this similarity, we computed the coefficient of determination (*R*^2^) between time-scaled trajectories for movements of the same direction but different durations (Fig. 4C); we use the coefficient of determination as it captures agreement in both the magnitude and the variation of the two trajectories (Methods). For every population, we found that the mean *R*^2^ between time-scaled trajectories corresponding to movements in the same direction was high (mean *R*^2^ = 0.64). It was significantly higher than our estimated lower limit for *R*^2^, given by the mean *R*^2^ between unscaled trajectories corresponding to arm movements of the same duration but in different directions (mean *R*^2^ = −>0.33; *t*-test for each population, max p-value *<* 10*^−^*^6^, *N* (same direction) = 48, *N* (different directions) = 336). We also estimated an upper limit for *R*^2^ (mean *R*^2^ = 0.72 over all populations) by comparing neural trajectories from movements of the same duration *and* direction (black bars in Fig. 4C; Methods). For 7 of the 13 populations there was no significant difference between this upper limit and the *R*^2^ between time-scaled trajectories (two-tailed *t*-test for each population: non-significant p-values 0.16 −> 0.7, M1=3, PMd=4; significant p-values 10*^−^*^6^ −> 0.04, M1=4, PMd=2). Across all populations, the time-scaled trajectories accounted for at least 92% of the difference between our estimated lower and upper limits for *R*^2^ (Fig. 4C). Consequently, nearly all the variation in neural dynamics between different durations of an arm movement in the same direction could be accounted for by temporal rescaling of a single, fixed neural trajectory.

To directly contrast the temporal scaling and geometry hypotheses, we tested whether the neural trajectories for different durations of arm movement were detectably separated. We found the distance between trajectories corresponding to different durations to be significantly smaller than the distances between trajectories from adjacent direction bins (Fig. 4D, *t*-test for each population, max p-value=3 *·* 10*^−^*^3^, *N* = 192), indicating that changes in movement duration had little influence on the magnitude of neural trajectories in M1 and PMd. As a more stringent control, we quantified the similarity of the neural trajectories for movements of the same duration *and* direction (within-bin trajectories: see Methods). We found that neural trajectories for the same duration and direction were as close together as scaled trajectories for different durations of the same direction of arm movement (Fig. 4E, one-sided *t*-test for each population, average p-value=0.8, *N* = 96). This result shows that neural trajectories for different durations of arm movement in the same direction are as close as they can be considering the natural variability of the neural activity. Collectively, these results support the temporal scaling hypothesis, that how quickly the arm moves is encoded by the rate at which population activity traverses a fixed trajectory (Fig. 4A, right).

### Maximum movement speed also has minimal effects on neural trajectories

We further tested the robustness of evidence for this hypothesis by repeating the distance analysis using maximum movement speed instead of duration as the temporal parameter of movement. To reduce the confounding effect of the scaling of maximum speed with distance (Fig. 1D), we selected a subset of movements for which distance was within a narrow range (Methods), so held approximately constant, but which resulted in considerably fewer movement samples (reduction of 23% on average across populations). Binning the neural activity into the same 8 direction bins but just high and low maximum speeds (averages 14 [cm/s] and 19 [cm/s] respectively) maximised the sample of movements to contrast at each speed to counteract the fewer samples overall. We obtained neural trajectories for each speed and direction bin by projecting the condition-averaged neural activity into the *D*-dimensional subspace defined above (2 speeds × 8 directions = 16 conditions total) and compared the distances between trajectories as in Fig. 4F. For all populations, the distance between trajectories corresponding to different maximum speeds was significantly smaller than the distance between trajectories corresponding to adjacent directions (*t*-test for each population, max p-value=3 *·* 10*^−^*^6^, *N* = 32). Despite its drawback of more strongly correlating with the spatial parameter of movement, using maximum movement speed to separate neural trajectories also supports a model in which movement speed is not encoded by changes in a neural trajectory but by how quickly it is traversed.

### Temporal rescaling of neural trajectories accurately decodes movement duration

To further test the temporal scaling hypothesis, we designed a decoder that explicitly assumes movement speed is encoded by how quickly a neural trajectory is traversed. The decoder estimated the duration of a sample trajectory (red in Fig. 5A, left) by comparing it to a reference trajectory (black in Fig. 5A, left) for the same movement direction. If the temporal scaling hypothesis were true then the sample and reference trajectories would be identical and the time at which the sample trajectory reached a particular point in state space would lead (if faster) or lag (if slower) the reference trajectory arriving at the same point (Fig. 5A illustrates leading). By assuming the sample and reference trajectories were identical, our decoder used the delays between them arriving at the same points to estimate the constant relative speed at which the sample trajectory was traversed, then used that speed to predict how long the sample trajectory and corresponding movement would be (Fig. 5A and legend). If the sample and reference trajectories were not the identical, then they need not arrive at the same point in state space, and need have no consistent lead or lag between the two: consequently our simple linear decoder would perform poorly.

**Figure 5:**
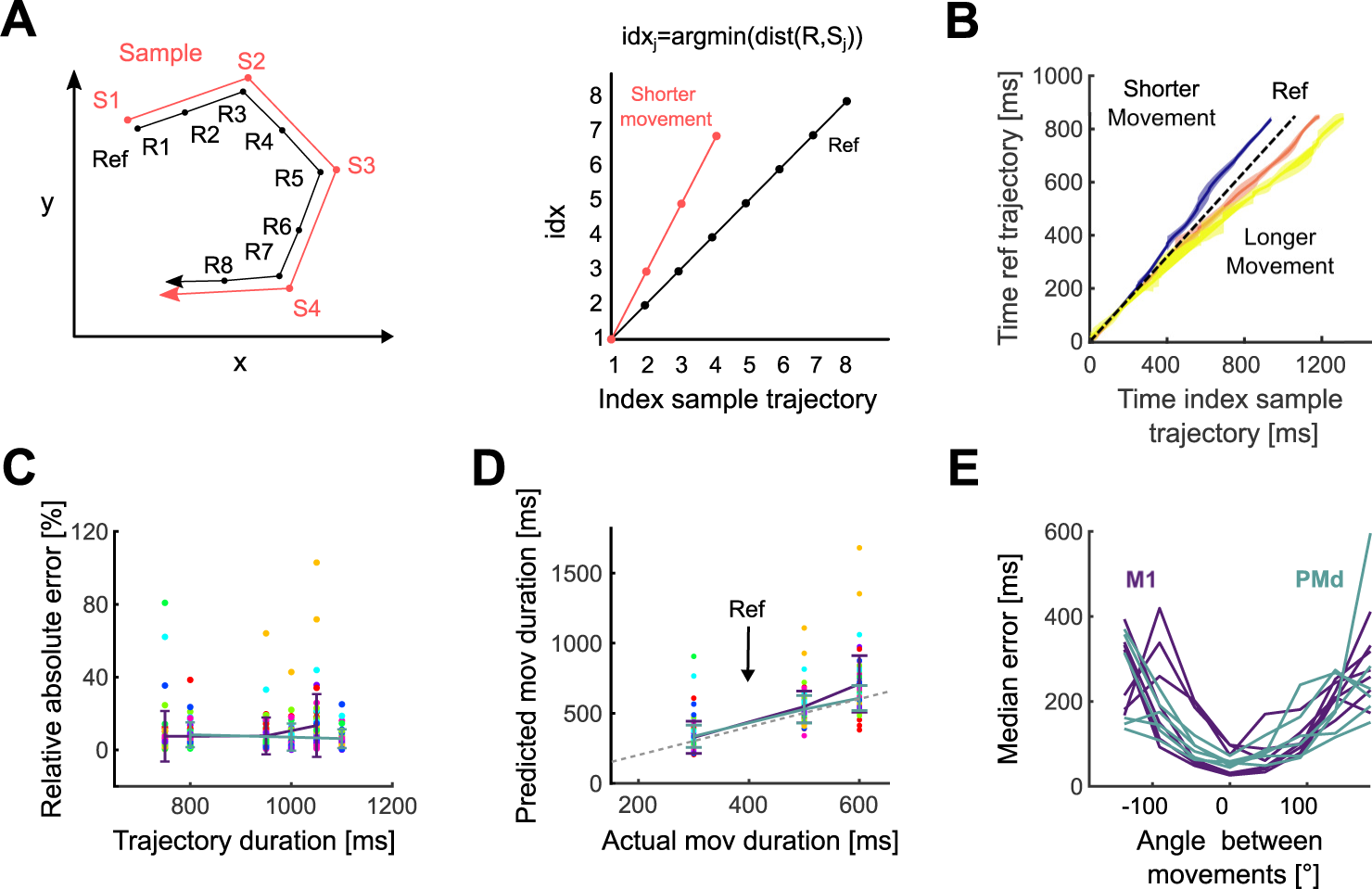
Movement duration can be decoded from the traversal of a fixed neural trajectory. A) Schematic of the duration decoder. Left: a sample trajectory compared to a reference trajectory for the same direction of movement. Right: time indices at which each point on the sample trajectory is closest to the reference trajectory. We estimated the relative speed (*v*) of the sample trajectory compared to the reference trajectory as the slope of a linear fit to the time index plot, and decoded *Duration_sample_* = *vDuration_ref_*. B) Time indices for sample trajectories from an example population. Using movements within the 300-400 ms duration bin as a reference (black), we calculated time indices for movements in the other three duration bins. Each coloured line shows the mean time index and standard deviation (across directions) for a given movement duration. C) Relative error of decoder predictions for trajectory duration. D) Predicted versus actual movement duration for all recordings. Coloured error bars indicate mean and SD for M1 and PMd (key in panel E). Coloured circles indicate different movement directions. The dashed line indicates the actual value of movement duration. E) Decoder error in predicting trajectory duration for reference and sample trajectories corresponding to different movement directions. One line per population.

For each population, we observed that sample neural trajectories consistently led or lagged the reference trajectory of 300-400 ms duration movements throughout a movement (Fig. 5B). Consequently, our linear decoder accurately predicted both the duration of neural trajectories across all populations (average relative error: 9.5% in M1 and 7.2% in PMd 5C) and the corresponding movement durations (Fig. 5D).

To verify that our decoder indeed performed poorly when the temporal scaling hypothesis was not true, we tested our decoder using a reference trajectory from a different direction of movement to that of the sample trajectory. The error in predicting trajectory durations increased with the difference between the movement directions corresponding to the reference and sample trajectories, and was up to two orders of magnitude larger than when using reference and sample trajectories from the same direction of movement (Fig. 5E). Our decoder thus provides further evidence that how quickly the arm moves is encoded by the rate at which population activity traverses a fixed trajectory through state space.

### Network modelling shows independent control of trajectories and their traversal

Finally, we used a recurrent neural network (RNN) to address the question of how independent parameters of a cortical network would allow independent control of the direction and speed of an arm movement. One hypothesis is that both direction and speed would be controlled by inputs to the network; indeed it has recently been argued that altering trajectory speed in a RNN, and hence altering arm movement speed, requires moving its dynamics to a different region of state-space (Saxena et al., 2022) However, our results here show that the neural trajectories for arm movements of different speeds are approximately the same, occupying the same region of state-space. We thus hypothesized that arm direction and speed are specified by different network parameters: the different neural trajectories for different movement directions specified by network inputs; and the traversal of trajectories specified by network parameter(s) that define the responsiveness of the network to the same input.

To test this, we built a recurrent neural network (RNN) based on a stability-optimised circuit model (Hennequin et al., 2014), composed of excitatory and inhibitory neurons whose synaptic connection strengths are optimised to produce a balanced, non-chaotic network (Methods). These networks are not trained to reproduce specific patterns of recorded neural activity yet can reproduce fundamental features of motor cortex dynamics at single-cell and population levels, such as the multi-phasic firing rate responses of individual units and the rotational trajectories in their low-dimensional subspace (Churchland et al., 2012; Hennequin et al., 2014).

Following Hennequin et al. (Hennequin et al., 2014), we modelled the input to the network during movement preparation as a ramp followed by a rapid decay. The input was projected into the *n*-neuron network by an *n*-dimensional weight vector randomly chosen from a uniform distribution ([−>1, 1]). Each neuron thus received a weighted, and possibly inverted, version of the same putative command signal for arm movement.

We first tested the hypothesis that the same input command signal could generate different trajectories of neural activity by rotating the *n*-dimensional vector that projected the input on to the network (Fig. 6A, see Methods). In this model, the angle of rotation between two projection vectors corresponds to the difference in the directions of the two required arm movements. Simulating this model with 10 rotations spanning 360*^◦^* gave low-dimensional neural trajectories that separated and reconverged (Fig. 6B and Fig. 6C), just as we observed in M1 and PMd (Fig. 3). Crucially, the distance between each pair of simulated trajectories was proportional to the angle between their input projection vectors (Fig. 6D), replicating how the distance between neural trajectories in motor cortex was proportional to the difference in their corresponding direction of arm movement (Fig. 3B). We found similar results for input projection vectors restricted to the range [0, 1] (results not shown), modelling solely excitatory inputs. Consequently, the separation of neural trajectories observed in motor cortex can be induced by rotating the projection of a common input signal.

**Figure 6:**
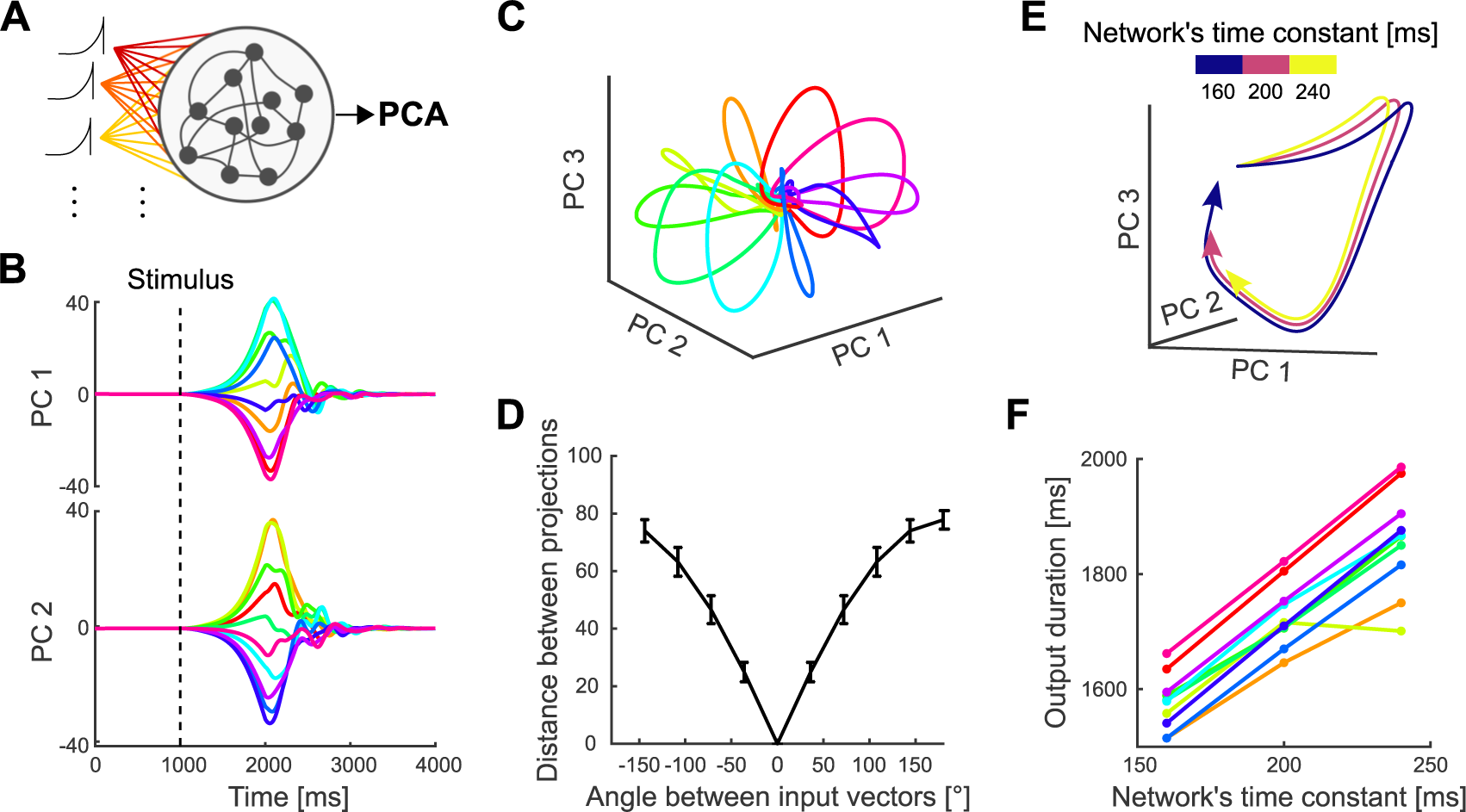
Independent control of movement direction and duration in a recurrent neural network (RNN). A) Schematic of an RNN whose input projection is rotated (colours) to simulate different movement directions. B) Trajectories of network activity in its PCA-defined subspace. Each colour indicates a rotation of the input projection. Dashed line shows the onset of input. C) Neural trajectories of an example network for each of the ten rotations of the input projection, simulating ten different directions of arm movement within 360 degrees. D) Average Euclidean distance between the network’s neural trajectories for different simulated arm movement directions. Error bars show standard deviation. E) First 1.9 s of the RNN’s neural trajectories for a fixed input projection and differing network time constants. F) Estimated duration of the neural trajectories for different time constants, one line per simulated movement direction. Duration was estimated by when the trajectory returned to its starting point in state space (panel C).

We then tested the hypothesis that the rate of traversing a trajectory could be specified by a parameter that controls the responsiveness of a network to the same input. We here focused on changing the network’s time constant parameter, which determines how quickly the neurons integrate their input (Eq.2, Methods; other parameters are considered in the Discussion). Simulating the network using a single input projection while only varying the time constant of the network, we found that trajectories generated with larger time constants were slower but with preserved shape (Fig. 6E), replicating the neural trajectories corresponding to different movement durations that we observed in motor cortex (Fig. 4B). This slower rate translated into trajectories of longer duration (Fig. 6F). Consequently, the RNN modelling shows that network parameters can independently specify the generation and traversal of neural trajectories, and so in principle independently control arm movement direction and speed.

## Discussion

The dynamical systems view of motor cortex proposes that its low-dimensional dynamics generate movement (Churchland et al., 2010, 2012; Shenoy et al., 2013; Michaels et al., 2016; Gallego et al., 2017; Russo et al., 2018). This view leaves two open questions: of how changes in those latent dynamics map to changes in the parameters of movement (Russo et al., 2018; Humphries, 2021b; Saxena et al., 2022); and of how those latent dynamics might be controlled to cause the change of their corresponding parameters of movement. Here we consider the case of independent spatial (direction) and temporal (duration) parameters of arm movement. While prior work has shown that direction is reflected in the geometry of the neural activity (Santhanam et al., 2009; Churchland et al., 2012; Michaels et al., 2016; Russo et al., 2018; Even-Chen et al., 2019), there is no clear hypothesis of how spatial and temporal parameters can be encoded and controlled simultaneously in the latent dynamics of motor cortex.

To examine how independent parameters of movement map on to changes in the latent dynamics, we took advantage of the variable direction and duration of arm movements naturally occurring in a sequential random-target task. This allowed us to ask whether parameters that can be specified independently in behaviour are also encoded independently in the dynamics of motor cortex. Studies of other cortical regions have found that, at a population level, task variables can be encoded in separate neural subspaces. This allows a separate, and sometimes orthogonal, representation of multiple variables in the same population (Kobak et al., 2016; Kaufman et al., 2016; Aoi et al., 2020; Saxena et al., 2022; Maggi and Humphries, 2022). Because of this, we explicitly tested for separate subspaces of activity encoding movement parameters. However, we found no evidence of a specific subspace related to movement duration, suggesting that the direction of an arm movement and how quickly it is made are not represented in distinct subspaces (Figure 2). Instead, we found that the direction and speed of arm movements respectively mapped to the direction and rate of traversal of low-dimensional trajectories of motor cortex activity. This mapping replicated across every recorded neural population, and regardless of whether they came from PMd or M1.

Our observation of fixed neural trajectories that preserved their shape across variations in how quickly the arm moved (Fig. 4) are at odds with a recent report of arm movement speed correlating with different trajectories of neural activity (Saxena et al., 2022). This difference could be explained by the selection of the parameters examined in each analysis: Saxena et al. (2022) analysed the effects of the speed of a cyclical arm movement separately for each direction of movement, so isolating the effects of speed on the trajectory of neural activity. We examined both the direction and duration of an arm movement together and found that direction had a much larger effect on the neural trajectory: indeed, we found how quickly the arm moved explained very little of the variation in neural activity (Fig. 2 and Fig. 4). So while we cannot say that the speed of movement had no effect on the trajectory of neural activity, our results suggest that the difference in the trajectory of neural activity between any two movements is dominated by differences in their direction rather than their speed: to that extent that a decoder assuming identical trajectories for different speeds of movement could well predict a movement’s duration. Alternatively, the point-to-point arm movements of the target task analysed here and the repeated rotational movements of the cycling task used by Saxena et al. (2022) could be encoded differently within motor cortex, perhaps within different subspaces (DePasquale et al., 2023), such that the speed of rotation in the cycling task does indeed notably alter the trajectory of neural activity.

Temporal scaling of neural activity has been previously observed in prefrontal cortex (Wang et al., 2018; Remington et al., 2018) and in the striatum (Gouv^ea et al., 2015) when animals are required to judge the duration of time intervals. However, in the same setting, temporal scaling is not present in the thalamus (Wang et al., 2018), and so is not an inevitable property of neural dynamics in a time-judgement task. Conversely, as the sequential-target task analysed here presented minimal time constraints, our results also show that temporal scaling of neural activity can be observed in the cortex even when there is no explicit timing required by a task.

To address the second open question of how low-dimensional dynamics in motor cortex might be controlled to alter the corresponding parameters of movement, we showed that the generation and traversal of neural trajectories can be independently controlled in a recurrent neural network model. In our simulations, the trajectories’ generation and traversal were changed by respectively rotating the vector of inputs and scaling the network’s time constant. These two mechanisms could be implemented in various ways in motor cortex. A recent study showed that the dynamics of motor cortex during planning and execution of movements is highly modulated by external inputs (Sauerbrei et al., 2020). This suggests a rotating input could be implemented by projections from either other cortical areas or the thalamus (Inagaki et al., 2022; Moll et al., 2023). The temporal scaling of neural trajectories, characterised here as a change in the time constant of the network, could also be approximated by modulating the neuronal gain across the population (Stroud et al., 2018). Global gain modulation could occur via neuromodulators (Thurley et al., 2008; Wei et al., 2014; Da Silva et al., 2018); gain modulation could also occur by changing tonic inputs to the network that drive the neurons to their saturating nonlinearity (Remington et al., 2018; Wang et al., 2018). Other network-level mechanisms for controlling the generation and traversal of neural trajectories may exist: our goal here was to show proof of principle that independent network-level mechanisms exist to control both observed features in the data.

From our data and modelling results arise three hypotheses for how low-dimensional neural trajectories in motor cortex activity can be manipulated to control multiple arm movement parameters simultaneously and independently. First, that the direction of the low-dimensional trajectory of neural activity controls the planar direction of arm movement. Second, that the rate of traversing a specific low-dimensional trajectory of neural activity controls how quickly the arm is moved in that direction. Third, that this separation of encoding allows spatial and temporal components of arm movements to be independently specified, as we see in behavioural tasks (Soechting and Lacquaniti, 1981; Nishikawa et al., 1999; Mazzoni et al., 2007), by network-level changes that can act on the time-scale of individual movements.

Further exploration of these ideas could usefully look at the relationship between low-dimensional dynamics and the other crucial parameter of arm movement to a target, its extent. One possibility is that motor cortex encodes the target’s position in coordinates centered on the current hand position, from which both direction and extent could be computed. However, behavioural data have long suggested the hypothesis that the direction and extent of arm movements are also independently specified (Ghez et al., 1991; Gordon et al., 1992; Ghez and Krakauer, 2000). This hypothesis arose from behavioural studies showing that the distributions of target errors in arm movements were independent between the direction and extent of movement (Favilla et al., 1990; Gordon et al., 1994; Favilla and De Cecco, 1996; Vindras et al., 2005), and that adapting direction and extent to new targets occurred at independent rates (Krakauer et al., 2000). Recent work from Dudman and colleagues (Park et al., 2022) has added further support for this hypothesis by showing that the extent and direction of limb movement are preferentially encoded by different populations of layer 5 neurons of murine motor cortex, and manipulating those populations independently affected extent. A fuller model of the mapping between low-dimensional dynamics of motor cortex and changes in movement parameters will thus require determining if and how extent is so encoded.

A fuller model would also require further understanding the implications of our results for when movement parameters are specified. An influential version of the dynamical systems hypothesis proposes that neural activity during preparation defines the initial conditions of a neural dynamical system that is largely autonomous during movement itself (Churchland et al., 2010; Shenoy et al., 2013; Kaufman et al., 2014; Elsayed et al., 2016; Even-Chen et al., 2019). In contrast to this hypothesis we have shown that how quickly the arm moves is coded by how quickly a fixed trajectory of activity is traversed, and that fixed trajectory begins before movement onset. Consequently, the temporal component of a movement need not be coded within the initial conditions at the time of movement onset.

## Competing interests

The authors declare no competing interests.

## Acknowledgements

This work was supported by the Medical Research Council (MRC) [grant numbers MR/J008648/1, MR/S025944/1] to M.D.H. Collection of the data was supported by the NIH [grant numbers F31 NS092356/ T32 HD07418] to M.G.P. and [grant number R01 NS074044] to L.E.M.

## Data availability statement

All data used to produce the figures in this paper (electrophysiological recordings and simultaneous behaviour) is available at https://doi.org/10.80507/dandi.123456/0.123456.1234.

## Code availability

All code used to produce the figures in this paper was developed in MATLAB and is available in our GitHub repository https://github.com/Humphries-Lab/Motor_Cortex_Simultaneous_Coding.

## Author contributions

Andrea Colins Rodriguez: Formal Analysis, Methodology, Software, Writing – Original Draft Preparation, Visualisation, Validation, Data Curation.

Matthew G. Perich: Investigation, Funding Acquisition, Writing – Review and Editing. Lee Miller: Funding Acquisition, Writing – Review and Editing.

Mark Humphries: Conceptualization, Methodology, Formal Analysis, Software, Writing – Original Draft Preparation, Supervision, Project Administration, Funding Acquisition.

## Notes

### Competing Interest Statement

The authors have declared no competing interest.

### Summary of Updates

New analyses added (Figure 2, dPCA)

## References

Aoi, M. C., Mante, V., and Pillow, J. W. (2020). Prefrontal cortex exhibits multidimensional dynamic encoding during decision-making. Nature neuroscience, 23(11):1410– 1420.

Ashe, J. and Georgopoulos, A. P. (1994). Movement parameters and neural activity in motor cortex and area 5. Cerebral cortex, 4(6):590–600.

Churchland, M. M., Cunningham, J. P., Kaufman, M. T., Foster, J. D., Nuyujukian, P., Ryu, S. I., Shenoy, K. V., and Shenoy, K. V. (2012). Neural population dynamics during reaching. Nature, 487(7405):51–56.

Churchland, M. M., Cunningham, J. P., Kaufman, M. T., Ryu, S. I., and Shenoy, K. V. (2010). Cortical preparatory activity: representation of movement or first cog in a dynamical machine? Neuron, 68(3):387–400.

Churchland, M. M., Santhanam, G., and Shenoy, K. V. (2006). Preparatory activity in premotor and motor cortex reflects the speed of the upcoming reach. Journal of Neurophysiology, 96(6):3130–3146.

Colins Rodriguez, A. (2021). Monkey (arm movement). Zenodo.

Da Silva, J. A., Tecuapetla, F., Paixão, V., and Costa, R. M. (2018). Dopamine neuron activity before action initiation gates and invigorates future movements. Nature, 554(7691):244–248.

DePasquale, B., Sussillo, D., Abbott, L., and Churchland, M. M. (2023). The centrality of population-level factors to network computation is demonstrated by a versatile approach for training spiking networks. Neuron.

Elsayed, G. F., Lara, A. H., Kaufman, M. T., Churchland, M. M., and Cunningham, J. P. (2016). Reorganization between preparatory and movement population responses in motor cortex. Nat Commun, 7:13239.

Even-Chen, N., Sheffer, B., Vyas, S., Ryu, S. I., and Shenoy, K. V. (2019). Structure and variability of delay activity in premotor cortex. PLoS Computational Biology, 15(2):1– 17.

Favilla, M. and De Cecco, E. (1996). Parallel direction and extent specification of planar reaching arm movements in humans. Neuropsychologia, 34:609–613.

Favilla, M., Gordon, J., Hening, W., and Ghez, C. (1990). Trajectory control in targeted force impulses. vii. independent setting of amplitude and direction in response preparation. Experimental brain research, 79:530–538.

Gallego, J. A., Perich, M. G., Miller, L. E., and Solla, S. A. (2017). Neural manifolds for the control of movement. Neuron, 94:978–984.

Gallego, J. A., Perich, M. G., Naufel, S. N., Ethier, C., Solla, S. A., and Miller, L. E. (2018). Cortical population activity within a preserved neural manifold underlies multiple motor behaviors. Nature communications, 9:4233.

Georgopoulos, A. P., Kalaska, J. F., Caminiti, R., and Massey, J. T. (1982). On the relations between the direction of two-dimensional arm movements and cell discharge in primate motor cortex. Journal of Neuroscience, 2(11):1527–1537.

Georgopoulos, A. P., Schwartz, A. B., and Kettner, R. E. (1986). Neuronal population coding of movement direction. Science, 233(4771):1416–1419.

Ghez, C., Hening, W., and Gordon, J. (1991). Organization of voluntary movement. Current opinion in neurobiology, 1:664–671.

Ghez, C. and Krakauer, J. (2000). The organization of movement. In Kandel, E., Schwartz, J., and Jessell, T., editors, Principles of Neural Science 4th Edition, chapter 33, pages 653–673. McGraw-Hill.

Glaser, J. I., Perich, M. G., Ramkumar, P., Miller, L. E., and Kording, K. P. (2018). Population coding of conditional probability distributions in dorsal premotor cortex. Nature Communications, 9(1):1–14.

Gordon, J., Ghilardi, M., and Ghez, C. (1992). Parallel processing of direction and extent in reaching movements. IEEE Engineering in Medicine and Biology Magazine, 11(4):92– 93.

Gordon, J., Ghilardi, M. F., and Ghez, C. (1994). Accuracy of planar reaching movements. i. independence of direction and extent variability. Experimental brain research, 99:97– 111.

Gouvêa, T. S., Monteiro, T., Motiwala, A., Soares, S., Machens, C., and Paton, J. J. (2015). Striatal dynamics explain duration judgments. eLife, 4:1–14.

Hennequin, G., Vogels, T. P., and Gerstner, W. (2014). Optimal control of transient dynamics in balanced networks supports generation of complex movements. Neuron, 82(6):1394–1406.

Humphries, M. D. (2021a). Strong and weak principles of neural dimension reduction. *Neurons, Behaviour*, Data analysis & Theory, 5.

Humphries, M. D. (2021b). Strong and weak principles of neural dimension reduction. *Neurons, Behavior*, Data analysis, and Theory, 5(2).

Inagaki, H. K., Chen, S., Ridder, M. C., Sah, P., Li, N., Yang, Z., Hasanbegovic, H., Gao, Z., Gerfen, C. R., and Svoboda, K. (2022). A midbrain-thalamus-cortex circuit reorganizes cortical dynamics to initiate movement. Cell, 185(6):1065–1081.e23.

Inoue, Y., Mao, H., Suway, S. B., Orellana, J., and Schwartz, A. B. (2018). Decoding arm speed during reaching. Nature Communications, 9(1):1–14.

Kaufman, M. T., Churchland, M. M., Ryu, S. I., and Shenoy, K. V. (2014). Cortical activity in the null space: permitting preparation without movement. Nature neuroscience, 17:440–448.

Kaufman, M. T., Seely, J. S., Sussillo, D., Ryu, S. I., Shenoy, K. V., and Churchland, M. M. (2016). The largest response component in the motor cortex reflects movement timing but not movement type. eneuro, 3(4).

Kobak, D., Brendel, W., Constantinidis, C., Feierstein, C. E., Kepecs, A., Mainen, Z. F., Qi, X.-L., Romo, R., Uchida, N., and Machens, C. K. (2016). Demixed principal component analysis of neural population data. elife, 5:e10989.

Krakauer, J. W., Pine, Z. M., Ghilardi, M. F., and Ghez, C. (2000). Learning of visuomotor transformations for vectorial planning of reaching trajectories. J Neurosci, 20:8916– 8924.

Lawlor, P. N., Perich, M. G., Miller, L. E., and Kording, K. P. (2018). Linear-nonlineartime-warp-poisson models of neural activity. Journal of Computational Neuroscience, 45(3):173–191.

Maggi, S. and Humphries, M. D. (2022). Activity subspaces in medial prefrontal cortex distinguish states of the world. Journal of Neuroscience, 42(20):4131–4146.

Mazzoni, P., Hristova, A., and Krakauer, J. W. (2007). Why don’t we move faster? parkinson’s disease, movement vigor, and implicit motivation. J Neurosci, 27:7105– 7116.

Michaels, J. A., Dann, B., and Scherberger, H. (2016). Neural Population Dynamics during Reaching Are Better Explained by a Dynamical System than Representational Tuning. PLoS Computational Biology, 12(11):1–22.

Moll, F. W., Kranz, D., Asensio, A. C., Elmaleh, M., Ackert-Smith, L. A., and Long, M. A. (2023). Thalamus drives vocal onsets in the zebra finch courtship song. Nature, 616(7955):132–136.

Moran, D. W. and Schwartz, A. B. (1999). Motor cortical representation of speed and direction during reaching. Journal of neurophysiology, 82(5):2676–2692.

Nishikawa, K. C., Murray, S. T., and Flanders, M. (1999). Do arm postures vary with the speed of reaching? Journal of Neurophysiology, 81(5):2582–2586.

Paninski, L., Fellows, M. R., Hatsopoulos, N. G., and Donoghue, J. P. (2004). Spatiotemporal tuning of motor cortical neurons for hand position and velocity. Journal of neurophysiology, 91(1):515–532.

Park, J., Phillips, J. W., Guo, J.-Z., Martin, K. A., Hantman, A. W., and Dudman, J. T. (2022). Motor cortical output for skilled forelimb movement is selectively distributed across projection neuron classes. Science Advances, 8(10):eabj5167.

Perich, M., Lawlor, P., Kording, K., and Miller, L. (2018a). Extracellular neural recordings from macaque primary and dorsal premotor motor cortex during a sequential reaching task.

Perich, M. G., Gallego, J. A., and Miller, L. E. (2018b). A neural population mechanism for rapid learning. Neuron, 100:964–976.e7.

Remington, E. D., Narain, D., Hosseini, E. A., and Jazayeri, M. (2018). Flexible Sensorimotor Computations through Rapid Reconfiguration of Cortical Dynamics. Neuron, 98(5):1005–1019.e5.

Russo, A. A., Bittner, S. R., Perkins, S. M., Seely, J. S., London, B. M., Lara, A. H., Miri, A., Marshall, N. J., Kohn, A., Jessell, T. M., Abbott, L. F., Cunningham, J. P., and Churchland, M. M. (2018). Motor Cortex Embeds Muscle-like Commands in an Untangled Population Response. Neuron, 97(4):953–966.e1–e8.

Russo, A. A., Khajeh, R., Bittner, S. R., Perkins, S. M., Cunningham, J. P., Abbott, L. F., and Churchland, M. M. (2020). Neural Trajectories in the Supplementary Motor Area and Motor Cortex Exhibit Distinct Geometries, Compatible with Different Classes of Computation. Neuron, pages 1–14.

Santhanam, G., Yu, B. M., Gilja, V., Ryu, S. I., Afshar, A., Sahani, M., and Shenoy, K. V. (2009). Factor-analysis methods for higher-performance neural prostheses. Journal of Neurophysiology, 102(2):1315–1330.

Sauerbrei, B. A., Guo, J. Z., Cohen, J. D., Mischiati, M., Guo, W., Kabra, M., Verma, N., Mensh, B., Branson, K., and Hantman, A. W. (2020). Cortical pattern generation during dexterous movement is input-driven. Nature, 577(7790):386–391.

Saxena, S., Russo, A. A., Cunningham, J., and Churchland, M. M. (2022). Motor cortex activity across movement speeds is predicted by network-level strategies for generating muscle activity. eLife, 11:e67620.

Schroeder, K. E., Perkins, S. M., Wang, Q., and Churchland, M. M. (2022). Cortical Control of Virtual Self-Motion Using Task-Specific Subspaces. The Journal of Neuroscience, 42(2):220–239.

Schwartz, A. B. (1992). Motor cortical activity during drawing movements: single-unit activity during sinusoid tracing. Journal of neurophysiology, 68(2):528–541.

Schwartz, A. B. (1994). Direct cortical representation of drawing. Science, 265(5171):540– 542.

Shenoy, K. V., Sahani, M., and Churchland, M. M. (2013). Cortical control of arm movements: a dynamical systems perspective. Ann Rev Neurosci, 36:337–359.

Soechting, J. F. and Lacquaniti, F. (1981). Invariant characteristics of a pointing movement in man. J Neurosci, 1:710–720.

Stroud, J. P., Porter, M. A., Hennequin, G., and Vogels, T. P. (2018). Motor primitives in space and time via targeted gain modulation in cortical networks. Nature Neuroscience, 21(12):1774–1783.

Thura, D. and Cisek, P. (2014). Deliberation and commitment in the premotor and primary motor cortex during dynamic decision making. Neuron, 81(6):1401–1416.

Thurley, K., Senn, W., and Lüscher, H. R. (2008). Dopamine increases the gain of the input-output response of rat prefrontal pyramidal neurons. Journal of Neurophysiology, 99(6):2985–2997.

Vindras, P., Desmurget, M., and Viviani, P. (2005). Error parsing in visuomotor pointing reveals independent processing of amplitude and direction. J Neurophysiol, 94:1212– 1224.

Vyas, S., Golub, M. D., Sussillo, D., and Shenoy, K. V. (2020). Computation through neural population dynamics. Annual review of neuroscience, 43:249–275.

Wang, J., Narain, D., Hosseini, E. A., and Jazayeri, M. (2018). Flexible timing by temporal scaling of cortical responses. Nature Neuroscience, 21(1):102–112.

Wang, W., Chan, S. S., Heldman, D. A., and Moran, D. W. (2007). Motor cortical representation of position and velocity during reaching. Journal of Neurophysiology, 97(6):4258–4270.

Wei, K., Glaser, J. I., Deng, L., Thompson, C. K., Stevenson, I. H., Wang, Q., Hornby, T. G., Heckman, C. J., and Kording, K. P. (2014). Serotonin affects movement gain control in the spinal cord. Journal of Neuroscience, 34(38):12690–12700.

